# Molecular Genetic Analysis of Rbm45/Drbp1: Genomic Structure, Expression, and Evolution

**DOI:** 10.1101/274647

**Authors:** Lauren E. Price, Abigail B. Loewen Faul, Aleksandra Vuchkovska, Kevin J. Lopez, Katie M. Fast, Andrew G. Eck, David W. Hoferer, Jeffrey O. Henderson

## Abstract

RNA recognition motif-type RNA-binding domain containing proteins (RBDPs) participate in RNA metabolism including regulating mRNA stability, nuclear-cytoplasmic shuttling, and splicing. Rbm45 is an RBDP first cloned from rat brain and expressed spatiotemporally during rat neural development. More recently, RBM45 has been associated with pathological aggregates in the human neurological disorders amyotrophic lateral sclerosis, frontotemporal lobar degeneration, and Alzheimer’s. Rbm45 and the neural developmental protein musashi-1 are in the same family of RDBPs and have similar expression patterns. In contrast to Musashi-1, which is upregulated during colorectal carcinogenesis, we found no association of *RBM45* overexpression in human colon cancer tissue. In order to begin characterizing RNA-binding partners of Rbm45, we have successfully cloned and expressed human RBM45 in an Intein fusion-protein expression system. Furthermore, to gain a better understanding of the molecular genetics and evolution of Rbm45, we used an *in silico* approach to analyze the gene structure of the human and mouse Rbm45 homologues and explored the evolutionary conservation of Rbm45 in metazoans. Human *RBM45* and mouse *Rbm45* span ∽17 kb and 13 kb, respectively, and contain 10 exons, one of which is non-coding. Both genes have TATA-less promoters with an initiator and a GC-rich element. Downstream of exon 10, both homologues have canonical polyadenylation signals and an embryonic cytoplasmic polyadenylation element. Moreover, our data indicate Rbm45 is conserved across all metazoan taxa from sponges (phylum Porifera) to humans (phylum Chordata), portending a fundamental role in metazoan development.

## Introduction

RNA recognition motif-type RNA (RRM)-binding domain (RBD) containing proteins (RBDPs) have diverse cellular functions. RBDPs have been demonstrated to regulate the stability of target mRNAs (e.g. CUGBP2 [Mukhopadhyay *et al.*, 2003], HuR, [Abe *et al.*, 1996], and Elav [Joseph *et al*., 1998]), the translatability of target mRNAs (e.g. musashi-1 [Imai *et al.*, 2001; Sureban *et al*., 2008], Elav/Hu protein family [Antic *et al*., 1999; Chung *et al*., 1997; Okabe *et al.*, 2001; Kasashima *et al*., 1999; Kawagishi *et al*., 2013], and CUGBP2 [Anant *et al*., 2001; Mukhopadhyay, 2003; Sureban *et al.*, 2007]), or both, in various tissues including neural and digestive (e.g. CUGBP2; Anant *et al.*, 2004; Mukhopadhyay *et al.*, 2003). Besides the canonical function of RNA binding, several RBDPs have been identified as being multifunctional proteins participating in RNA-mediated protein:protein interactions (A1CF [Blanc *et al.*, 2001a, 2005, 2007; Galloway *et al.*, 2010; Henderson *et al.*, 2001; Mehta and Driscoll, 2002], Syncrip (GRY-RBP) [Blanc *et al.*, 2001b; Lau *et al.*, 2002], and CUGBP2 [Anant *et al.,* 2001; Mukhopadhyay *et al.*, 2003]) or RNA-independent protein:protein interactions (e.g. neuronal proteins TDP-43 and FUS, reviewed in Ugras and Shorter, 2012).

RNA-binding motif protein 45 (Rbm45), also known as developmentally regulated RNA-binding protein 1(Drbp1), was first identified as a novel RNA-binding protein through the screening of a rat cerebral expression library with polyclonal antibody against consensus RRM sequences. Rbm45 has three (Li *et al*., 2015b; Mashiko *et al*., 2016) or four canonical RBDs (Tamada *et al.*, 2002; HomoloGene, https://www.ncbi.nlm.nih.gov/homologene/?term=Rbm45 [accessed 11.15.2017]) and is expressed in a spatiotemporal manner in developing rat brain, with highest expression, as measured by mRNA accumulation, occurring between embryonic days 12 to 16. There is a subsequent decline in expression of Rbm45 during late embryonic development that extends into adulthood as neuronal precursor cells differentiate (Tamada *et al.*, 2002). Additionally, Rbm45 has been demonstrated to be upregulated after spinal cord injury in an opossum neonatal model of neuronal repair (Mladinic *et al*., 2010). Subsequent work from Robert Bowser’s lab group (2012, 2015a, 2015b, 2016) and Hitoshi Endo’s lab (2016) strongly associate RBM45 aggregation in the pathogenesis of the destructive neurodegenerative disorders amyotrophic lateral sclerosis (ALS) and frontotemporal lobar degeneration with TDP-43 inclusions (FTLD-TDP). Furthermore, mouse *Rbm*45 demonstrates strong association with aberrant prepulse inhibition, a measure of sensorimotor gating (startle reflex), an endophenotypic marker for neuropsychiatric disorders (Sittig *et al*., 2016). These studies implicate Rbm45 in neurogenesis, neuronal pathology, and neuronal repair. Despite the work of these groups, little is known about how Rbm45 accumulation contributes to the progression of neurodegenerative diseases, neural development, or neural maintenance.

RBDPs are typically multifunctional and act both in the nucleus and cytoplasm, influencing transcription, RNA splicing, RNA export, translation, and transport of mRNAs (reviewed in Ling *et al*., 2013). Current research on ALS and FTLD-TDP (Li *et al*., 2015b; Mashiko *et al*., 2016) indicates RBM45 shares some of these functions (e.g. cytoplasmic-nuclear shuttling, homo-oligomerization, and heterodimerization with proteins involved in splicing), but also interacts with other RNA-binding proteins to modulate responses to oxidative stress and extracellular signaling (Li *et al*., 2016). In this report, we analyze the genomic structure, regulatory elements, gene expression, and evolutionary history of Rbm45 to gain an understanding of its role in metazoan development.

## Experimental Procedures

### Genomic Analysis

Rbm45 nucleotide and protein sequences were obtained from the National Center for Biotechnology Information (NCBI; https://www.ncbi.nlm.nih.gov/). Rbm45 orthologues were identified using the BLAST algorithm with *Mus musculus* (mouse; strain: Balb / C) Rbm45 cDNA, accession number (no.) AB036992.1 (Tamada *et al.*, 2002) as the query sequence and retrieved from GenBank (https://www.ncbi.nlm.nih.gov/). The following accession nos. were utilized in this study: mouse (strain: C57BL/6J) Rbm45 genomic nucleotide (nt) sequence (NT039207.6), *Homo sapiens* (human) Rbm45 genomic nt sequence (NT005403), human Rbm45 cDNA (AB03699.1) and protein (NP_694453.2), mouse Rbm45 protein (AAH57890.1), *Rattus norvegicus* (Norway rat) Rbm45 cDNA (AB036990) and protein (NP_695218), *Pan troglodytes* (chimpanzee) Rbm45 cDNA (XM_515938.2) and protein (JAA06242), *Macaca mulatta* (Rhesus monkey) Rbm45 cDNA (XM_001097905.1) and protein (AFE64478), *Pongo abelii* (Sumatran orangutan) Rbm45 cDNA (CR857895.1) and protein (NP_001125039), *Monodelphis domestica* (gray short-tailed opossum) Rbm45 cDNA (XM_001368729.1) and protein (XP_007494492), *Canis lupus familiaris* (dog) Rbm45 cDNA (XM_535977.2) and protein (XP_013966212), *Gallus gallus* (chicken) Rbm45 cDNA (NM001030252.1) and protein (NP_001026423.1), *Xenopus laevis* (African clawed frog) Rbm45 cDNA (BC045039.1) and protein (NP_001080090.1), and *Danio rerio* (zebra fish) Rbm45 cDNA (XM_691856.2) and protein (NP_001120874.1).

All sequence alignments were performed using ClustalW (v1.83; http://www.ebi.ac.uk/clustalw/) or ClustalOmega (v1.2.1; https://www.ebi.ac.uk/Tools/msa/clustalo/). Molecular phylogenetic trees were created using PhyML3.1 (http://www.atgcmontpellier.fr/phyml/; Guindon *et al.*, 2010) driven by SeaView-Multiplatform GUI for Molecular Phylogeny v4.6.2 (http://doua.prabi.fr/software/seaview; Gouy *et al.*, 2010). Molecular phylogenetic tree creation with Rbm45 nucleotide sequences used the following parameters: Model: GTR; Branch Support: aLRT (SH-like); Nucleotide Equilibrium Frequencies: Empirical; Invariable Sites: None; Across Site Rate Variation: Optimized; Tree Searching Operations: NNI; Starting Tree: BioNJ Optimize tree topology. Molecular phylogenetic tree creation with Rbm45 amino acid sequences used the following parameters: Model: LG; Branch Support: aLRT (SH-like); Amino Acid Equilibrium Frequencies: Model-given; Invariable Sites: None; Across Site Rate Variation: Optimized; Tree Searching Operations: NNI; Starting Tree: BioNJ Optimize tree topology.

### Tissue Procurement

C57Bl / 6J male mice were obtained from The Jackson Laboratory. Mice were maintained and sacrificed as approved by the Animal Studies Committees of Washington University School of Medicine and Tabor College. Tissues were harvested from 5 week old mice, washed in ice-cold phosphate buffered saline (PBS; 10 mM Na_2_HPO_4_, 1.8 mM KH_2_PO_4_, 137 mM NaCl, 2.7 mM KCl, pH 7.4) and snap frozen using a dry ice / ethanol bath (– 75°C). Human normal adjacent and cancerous colon tissue was kindly provided by Dr. Nicholas O. Davidson (Washington University School of Medicine) and obtained from The Siteman Cancer Center Tissue Procurement Core under a protocol approved by the Washington University-St. Louis Human Research Protection Office.

### Quantitative Reverse Transcription-Polymerase Chain Reaction (qRT-PCR) Amplification Analysis of Human and Mouse Rbm45

RNA was extracted from normal adjacent and tumor tissue from human colon using TRI-reagent (Sigma-Aldrich) as described by the manufacturer. Total RNA for reverse transcription was prepared as described by Henderson and coworkers (2001), with the following modifications. Briefly, aliquots of total RNA (10 μg) were treated with 2 units RQ1 DNase (Promega) at 37°C for 1 h in 100 μL 40 mM Tris-HCl pH 7.6, 6 mM MgCl_2_, 10 mM NaCl, 40 U RNasin (Promega). The RNA was sequentially extracted with phenol:chloroform and chloroform, precipitated with ethanol at – 80°C, washed once with 75% ethanol and resuspended in diethylpyrocarbonate-treated water. RNA concentration was confirmed by measuring the absorbance at 260 nm. Five hundred nanograms of RNA were used for reverse transcription in the presence of 200 ng random hexamer (dN_6_) and 200 units SuperScript II RNase H^‒^ Reverse Transcriptase (Life Technologies) as directed by the manufacturer. qRT-PCR analysis of human and mouse Rbm45, and 18s rRNA was performed using an ABI Prism 7700 Sequence Detection System (Applied Biosystems) using SYBR green PCR master mix according to the manufacturer’s instructions (Applied Biosystems). Thermal cycling conditions were as follows: 50°C for 2 min, 95°C for 10 min, then 40 cycles of 95°C for 15 sec and 60°C for 1 min. Primers were designed using Primer Express v2.0.0 software (Applied Biosystems) and amplify coding DNA separated by at least one intron. Primers were compared to the public nucleotide database (https://www.ncbi.nlm.nih.gov/) by BLAST analysis to confirm specificity. Primers for human RBM45 are QhRBM45F (sense; 5′-cttccttgtgatcagcaagtacaca-3′) and QhRBM45R (antisense; 5′-gtcctggatgtcgccaaaa-3′). Primers for mouse Rbm45 are QmRbm45F (sense; 5′-tgttgtacttccgtcgtgcaa-3′) and QmRbm45R (antisense; 5′-ttaaacacaacaaacagcctttctttc-3′). Primers for 18s rRNA are 18sF (sense; 5′-cggctaccacatccaaggaa-3′) and 18sR (antisense; 5′-gctggaattaccgcggct-3′). The data were normalized to 18s rRNA levels in each sample, then 2^-ΔΔCt^ was calculated, as described by the manufacturer (User Bulletin #2 [10/2001], Applied Biosystems), to normalize mRNA abundance relative to normal tissue in the case of human colon cancer samples, or brain, in the analysis of adult mouse tissues. Statistical analysis was done using Microsoft Excel. Student’s *t* test was used to determine *P* values. 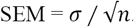.

### Cloning and expression of recombinant Rbm45

Human RBM45 cDNA was isolated from FirstChoice® Human Brain Total RNA (Ambion) using oligo(dT)_20_ and SuperScript III RNase H^‒^ Reverse Transcriptase (Invitrogen), as described by the manufacturer, and subsequently amplified with AccuPrime *Pfx* DNA polymerase (Invitrogen) in a Dyad thermal cycler (MJ Research) using primers hRBM45IntF (sense; 5′-ggt ggt cat atg gac gaa gct ggc agc tct-3′; *Nde*I restriction site underlined, amino acid [AA] 2 double underlined) and hRBM45IntR (antisense; 5′-ggt ggt tgc tct tcc gca gta agt tct ttg ccg ttt-3′; *Sap*I restriction site underlined, AA474 double underlined) under the following cycling conditions: Initial heat denaturation at 95°C for 2 min, then 35 cycles of 95°C for 15 sec, annealing gradient from 45°C to 65°C for 30 sec, and extension at 68°C for 1 min 30 sec. A final extension of 10 min at 72°C was used for the last cycle. The 1449 base pair (bp) amplicon produced from annealing at 56.9°C was subcloned into pCR-Blunt II-TOPO (Invitrogen), to give plasmid pCRII-hRBM45, and sequenced on both strands.

The human RBM45 cDNA was then cloned as an *Nde*I-*Sap*I fragment from pCRII-hRBM45 into pTXB1 (New England Biolabs) creating a human RBM45 carboxyl-terminus (C-terminus) Intein fusion protein. The Intein tag is self-cleavable and contains a chitin-binding domain for affinity purification as a fusion protein. Expression and purification of the fusion protein was conducted in ER2566 *E. coli* cells as previously described (Blanc *et al*., 2001b), with the exception that cells were induced with 0.4 mM isopropyl β-D-1-thiogalactopyranoside (IPTG), grown at 37°C, and lysed with B-PER Bacterial Protein Extraction Reagent (Pierce Biotechnology). Protein fractions were separated by 10-20% gradient SDS-PAGE and stained with Coomassie Brilliant Blue R-250 (BIO-RAD).

Mouse Rbm45 cDNA was cloned using a nested PCR approach. ssDNA was produced from mouse strain C57Bl/6J brain total RNA, purified with TRIzol Reagent (Invitrogen), using the gene specific primer mRbm45R (antisense; 5′-cagtcagtcagctgaggattatgtc-3′) and SuperScript III RNase H^‒^ Reverse Transcriptase (Invitrogen) as described by the manufacturer. The Rbm45 ssDNA was used as a template to amplify a full-length cDNA, flanked by non-coding DNA, using AccuPrime *Pfx* DNA polymerase (Invitrogen) and primers mRbm45F (sense; 5′-agaacctgccgggtgaaca-3′) and mRbm45R under the following cycling conditions: Initial heat denaturation at 95°C for 2 min, then 30 cycles of 95°C for 15 sec, annealing at 55°C for 30 sec, and extension at 68°C for 1 min 30 sec. A final extension of 10 min at 72°C was used for the last cycle. The 1542 bp amplicon was subcloned into pSC-B using the StrataClone Ultra Blunt PCR Cloning Kit (Agilent) to create pJH1 and sequenced on both strands. The mouse Rbm45 coding sequence was then amplified from pJH1 using primers mRbm45IntF (sense; 5′-ggt ggt cat atg gac gac gcc ggc ggc tt-3′; *Nde*I restriction site underlined, AA2 double underlined) and mRbm45IntR (antisense; 5′-gta agt tct ctg cct ctt ttt-3′; AA476 double underlined) under the above cycling conditions. The 1455 bp amplicon was cleaved with *Sap*I and cloned into pTYB2 (New England Biolabs) digested with *Nde*I and *Sma*I creating a mouse Rbm45 C-terminal Intein fusion protein. Expression and purification of the fusion protein was conducted in ER2566 *E. coli* cells as previously described (Blanc *et al*., 2001b) with the following modifications: Cells were grown at 15°C, 25°C, or 30°C; each temperature point was uninduced or induced with 0.3 mM, 0.4 mM, or 0.5 mM IPTG. Cells were lysed with B-PER Bacterial Protein Extraction Reagent (Pierce Biotechnology) and total protein lysate analyzed by 10-20% gradient SDS-PAGE stained with Coomassie Brilliant Blue R-250 (BIO-RAD).

## Results and Discussion

### Human RBM45: Genomic Organization and Structure

Using BLASTn (https://blast.ncbi.nlm.nih.gov), we queried (Spring 2007) the non-redundant nucleotide collection (nr/nt) with human cDNA sequence revealing the exon/intron junctions on the human genomic sequence. We used this information to build a map of the *RBM45* locus. The chromosomal locus containing the *Homo sapiens* (human) *RBM45* gene spans approximately 17 kb and contains 10 exons (Fig. 1A), matching NCBI annotation data (26-Jan-2014, NM_152945). Database analysis (http://www.ncbi.nlm.nih.gov/Uni-Gene/clust.cgi) localizes the gene (UniGene Hs.377257; UGID:201961) to chromosome 2q31.2 (Assembly: GRCh38.p7 [GCF_000001405.33]; Location: NC_000002.12 [178112409…178129656 nt]) and having a plus orientation. The *RBM45* gene is contained within chromosome 2 genomic scaffold GRCh38.p7 primary assembly HSCHR2_CTG7_2, accession no. NT_005403.18, and contains complementary exon sequence information to that obtained by our cloning and sequencing the human cDNA from brain (data not shown).

**Figure 1.**
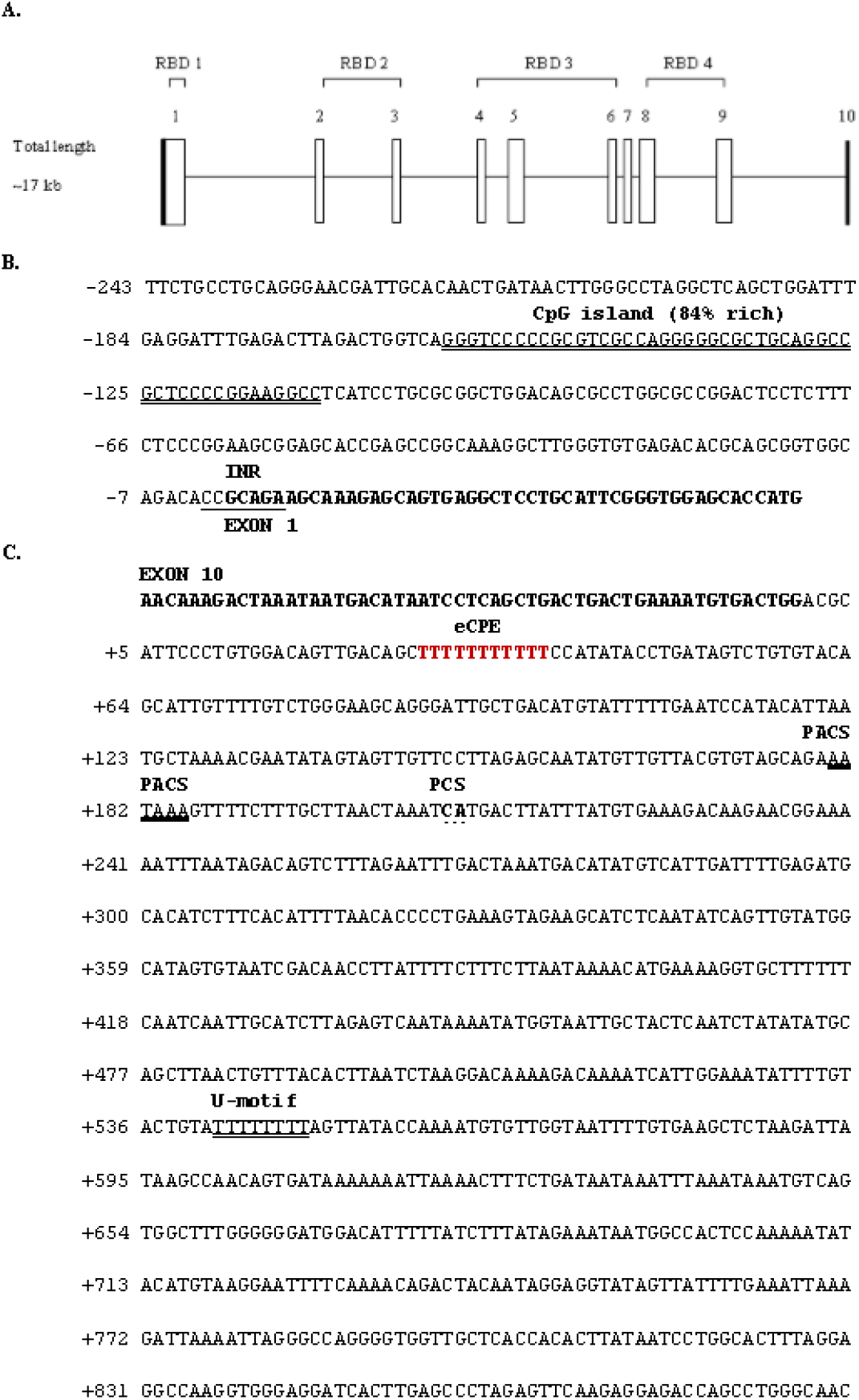
Genomic structure of the human*RBM45*gene. The human *RBM45* gene has 10 exons. (A) (Top) The exons containing the four RNA-binding domains (RBD) are indicated by brackets. (Bottom) Schematic diagram of the exon-intron structure of the human *RBM45* gene. Vertical boxes represent the non-coding (filled boxes) and coding exons (open boxes), and are numbered sequentially. Solid horizontal lines represent introns. The diagram reflects the relative sizes of the exons and introns and is not drawn completely to scale. The introns and exons are scaled by type; the intron width scale is half that of the exon width scale (B) A portion of exon 1 (bold type) is shown with 243 upstream nucleotides. The ATG start of translation is the last three nt. Putative promoter elements identified are an initiator element (INR), underlined, and a CpG island, double underlined. (C) Exon 10 (bold type) with 889 downstream nucleotides. Predicted polyadenylation signaling elements identified are an embryonic cytoplasmic polyadenylation element (eCPE), red lettering; a canonical polyadenylation consensus signal (PACS), thick underline; polyadenylation cleavage site (PCS), dotted underline; and a U-rich motif (U-motif), double underlined.

Further analysis of the NCBI database revealed an inferred processed-pseudogene (i.e. retropseudogene; NG_026133.1 derived from AC060788.11) on chromosome 8q24.22 (Assembly: GRCh38.p7 [GCF_000001405.33]; Location: NC_000008.11 [131712316…131712980]). Retropseudogenes arise from reverse transcription of the parental gene RNA with subsequent random genomic integration of the resulting cDNA. Because the cDNA lacks upstream promoter elements, it is typically not expressed unless integration arbitrarily occurs downstream of an existing promoter. In the case of integration downstream of a promoter element, with concomitant protein expression, the retropseudogene is classified as a retrogene (Strachan *et al.*, 2015). Currently (10.21.2017), no information exists as to whether the human *RBM45* pseudogene is a retrogene. Nevertheless, the four RBDs described in the rat Rbm45 cDNA cloned by Endo and colleagues (Tamada *et al*., 2002) are the products of exons 1, 2-3, 4-6, and 8-9, respectively (Fig. 1A).

Approximately 243 nt of sequence 5′ of exon 1 was analyzed for putative promoter elements. The core promoter was found to be TATA-less, a common occurrence in genes that regulate development in metazoans. Confounding the identification of potential promoter elements, both in human and mouse, is the observation that TATA-less promoters can potentially have many different start sites ranging from 20 to 200 bp upstream of the 5′-UTR (Weaver, 2012). Nevertheless, we identified putative core promoter elements for the *RBM45* gene including an initiator element (INR) conforming roughly to the canonical PyPyAN(T/A)PyPy (where Py is any pyrimidine, e.g. thymine or cytosine, and N is any nucleotide) which would be predicted to encompass the transcription start site. INR sequences are highly degenerate (Weaver, 2012) and our putative INR matches 4 of 7 of the canonical nucleotides. Confirming this region as the transcription start site would require performing primer extension analysis (Henderson *et al.*, 2001), for example, but is beyond the scope of this report. In addition to the INR, a CpG island, a proximal promoter element, was identified suggesting possible epigenetic regulation of *RBM45* via cytosine methylation (Deaton and Bird, 2011). Both elements have been reported (Weaver, 2012) to be important for transcription initiation in the absence of a TATA box. Interestingly, a downstream promoter element (DPE) was not identified in the human *RBM45* exon 1 5′-untranslated region (5′-UTR; Fig. 1B). DPEs are typically found + 28 to + 32 relative to the transcription start site and are a common feature in TATA-less promoters. In contrast, the related mouse *Rbm45* contains a DPE in its 5′-UTR (Fig. 2B). However, presence of a DPE in TATA-less promoters is not sacrosanct as eukaryotic promoters are highly modular (Weaver, 2012).

**Figure 2.**
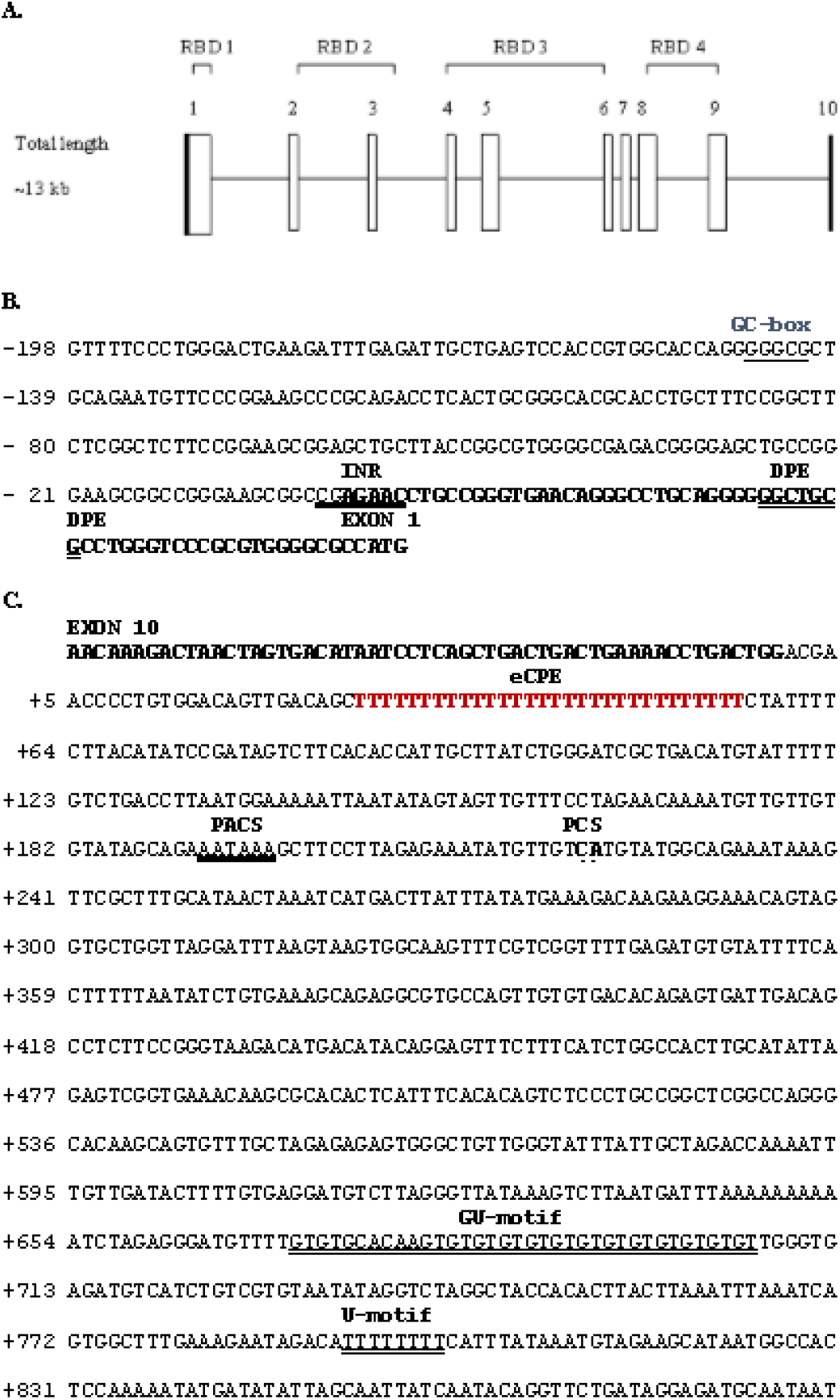
Structure of the mouse *Rbm45* gene. The mouse *Rbm45* gene has 10 exons. (A) (Top) The exons containing the four RNA-binding domains (RBD) are indicated by brackets. (Bottom) Schematic diagram of the exon-intron structure of the mouse *Rbm*45 gene. Vertical boxes represent the non-coding (filled boxes) and coding exons (open boxes), and are numbered sequentially. Solid horizontal lines represent introns. The diagram reflects the relative sizes of the exons and introns and is not drawn completely to scale. The introns and exons are scaled by type; the intron width scale is half that of the exon width scale (B) The sequences in bold represent the predicted 5′ UTR of exon 1 with 198 upstream nucleotides. The ATG start of translation is the last three nt. Putative promoter elements identified are a GC-box, thin underline; initiator element (INR), thick underline; and a downstream promoter element (DPE), double-underlined. (C) Exon 10 (bold type) with 889 downstream nucleotides. Predicted polyadenylation signaling elements identified are an embryonic cytoplasmic polyadenylation element (eCPE), red lettering; a canonical polyadenylation consensus signal (PACS), solid underlined; polyadenylation cleavage site (PCS), dotted underline; and a GU/U-rich motif, double-solid underlined.

Messenger RNA (mRNA) molecules often possess a poly(A) tail of approximately 250 adenosine residues. The poly(A) tail acts to increase the stability (Huez *et al.*, 1974) and translatability (Munroe and Jacobson, 1990) of mRNA; additionally, the poly(A) tail plays a role in the transport of mature mRNA molecules from the nucleus to the cytoplasm (Eckner *et al.*, 1991). Approximately 889 nt of genomic DNA (gDNA) downstream of *RBM45* exon 10 were examined for putative polyadenylation signal elements, revealing the presence of an embryonic cytoplasmic polyadenylation signal (Fig. 1C; Fox *et al*., 1989) upstream of a canonical polyadenylation consensus signal (PACS; Fitzgerald and Shenk, 1981). Downstream of the PACS is a polyadenylation cleavage site (PCS) and a U-rich motif (Fig. 1C). Typically, the U-rich motif (T-rich in gDNA) is immediately preceded by a GU-rich motif (GT-rich in gDNA) but is not required for polyadenylation; albeit, polyadenylation is suboptimal in the absence of a GU-rich motif (Gil and Proudfoot, 1987; Weill *et al*., 2012).

All of the *RBM45* intron-exon junctions were analyzed and conformed to the canonical consensus for splice donor and splice acceptor sites (Breathnach and Chambon, 1981) as enumerated in Table 1. Sequence analysis of cDNA clones from RT-PCR of adult human brain RNA (data not shown) did not reveal splice variants, consistent with previous analyses (Mammalian Gene Collection (MGC) Program Team, 2002). However, database interrogation (https://blast.ncbi.nlm.nih.gov/Blast.cgi) uncovered two predicted RBM45 variant, X1 (XP_016858809.1) and X2 (XP_005246344.1) derived by NCBI’s automated computational analysis using gene prediction method Gnomon (deposited 06.16.2016; https://www.ncbi.nlm.nih.gov/nuccore/1034611506/). Inspection of the cDNA sequence (XM_017003320.1) of the predicted RBM45 X1 isoform demonstrates it being derived through an alternative splice acceptor site adding 429 nt of immediately adjacent nt from the 5′-end of intron 1 (143 novel AA) to the 3′-end of exon 1 compared to wild-type RBM45. The resulting variant is predicted to encode a 617AA protein, compared to 474AA in the wild-type protein. Analysis of the cDNA sequence (XM_005246287.4) of the predicted RBM45 X2 isoform shows it is derived by use of a cryptic splice acceptor site on the 5′-end of exon 5, resulting in an in-frame 2 AA insertion predicted to encode a 476AA protein. Neither RBM45 isoform X1 or X2 is known to be physiologically relevant.

**Table 1.**
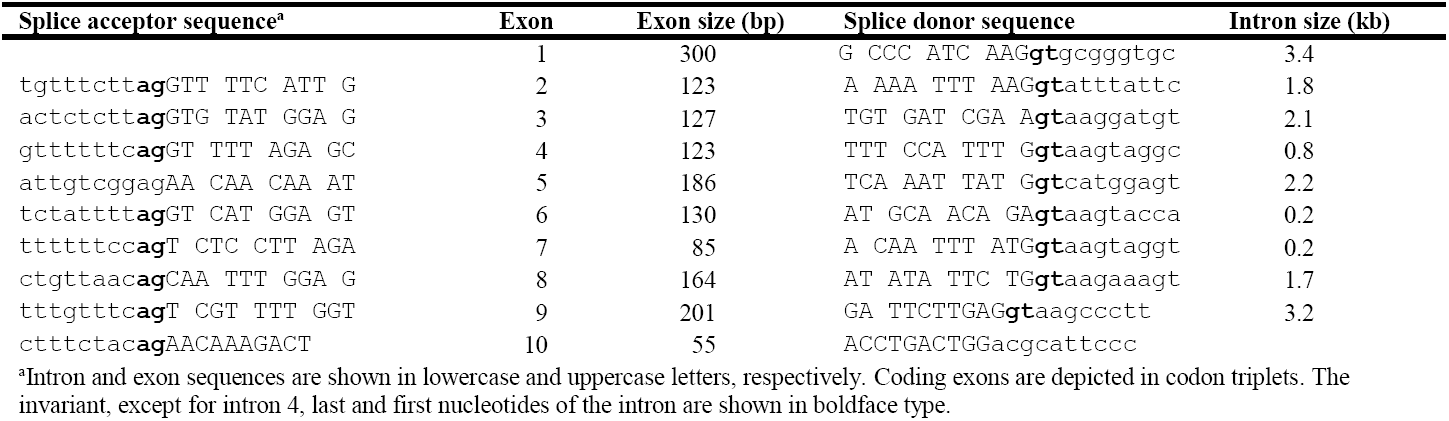
Exon-inron boundaries of human *RBM45*

### Mouse Rbm45: Genomic Organization and Structure

As with the human sequence, above, we queried (Spring 2007) the nr/nt collection, using BLASTn, with the mouse cDNA sequence revealing the exon/intron junctions on the mouse genomic sequence. Similar to human, the chromosomal locus containing the *Mus musculus* (mouse) *Rbm45* gene spans approximately 13 kb and contains 10 exons (Fig. 2A), matching NCBI annotation data (06-Oct-2010; AK080420.1, NM_153405.2). Data base analysis (http://www.ncbi.nlm.nih.gov/UniGene/clust.cgi) localizes the gene (UniGene Mm.33310; UGID:266774) to chromosome 2C3|2 (Assembly: GRCm38.p4 [GCF_000001635.24]; Location: NC_000068.7 [76369958…76383768 nt]). The *Rbm45* gene is contained within GenBank BAC clone RP24-456G2 (Clone_lib: RPCI-24), accession no. AC122048.2, and contains complementary exon sequence information obtained by our cloning and sequencing the mouse cDNA from brain (data not shown). The four RBDs described in the cDNA cloned by Endo and coworkers (Tamada *et al.*, 2002) are the products of exons 1, 2-3, 4-6, and 8-9, respectively (Fig. 2A).

Like human *RBM45*, analysis of mouse *Rbm45* sequence 5′ of exon 1 (approximately 198 nt) reveals that the promoter is TATA-less and contains a putative degenerate consensus (5 of 7 nt match) INR encompassing a predicted transcription start site (compare Fig. 2B to Fig. 1B). In contrast to human *RBM45* (Fig. 1B), mouse *Rbm45* contains a sequence with 4 of 7 nt matches to the consensus ([A/G]G[A/T]CGTG) DPE sequence at + 28 to + 32 relative to the predicted transcriptional start site, and has a GC-rich proximal promoter element in the form of a canonical GC-box (Fig. 2B). GC-boxes have been demonstrated to enhance transcription of TATA-less promoters by binding the transcription factor Sp1 (Goodrich *et al.*, 1996).

As in human *RBM45*, analysis of approximately 889 nt of gDNA downstream of mouse *Rbm45* exon 10 for putative polyadenylation signal elements (Fig. 2C) revealed an embryonic cytoplasmic polyadenylation consensus signal upstream of a PACS. Downstream of the PACS is a PCS and, in contrast to human *RBM45*, both a canonical GU-and U-rich motif. The presence of both of these elements suggests that mouse *Rbm45* may be more efficiently polyadenylated than human *RBM45*, thereby increasing expression of Rbm45 protein (Gil and Proudfoot, 1987). This assertion would need to be confirmed empirically, but is a reasonable hypothesis given the sequence data. In both the human and mouse Rbm45 genes, the presence of an embryonic cytoplasmic polyadenylation consensus signal correlates with the observation that highest expression of *RBM45*/*Rbm45* occurs during (neural) development, at least in the rat and mouse (Tamada *et al*., 2002; McKee *et al*., 2005). Analysis of human fetal brain tissue by qRT-PCR (Henderson *et al.*, 2001) would give some indication of the developmental regulation of human *RBM45* mRNA levels; unfortunately, such a study is beyond the capabilities of our laboratory.

All of the mouse *Rbm45* intron-exon junctions were analyzed and conformed to the canonical consensus splice donor and splice acceptor sites (Breathnach and Chambon, 1981) as enumerated in Table 2. Sequence analysis of cDNA clones from RT-PCR and database interrogation (https://blast.ncbi.nlm.nih.gov/Blast.cgi; data not shown) did not reveal splice variants, consistent with previous analyses (Tamada *et al*., 2002; McKee *et al*., 2005).

**Table 2.**
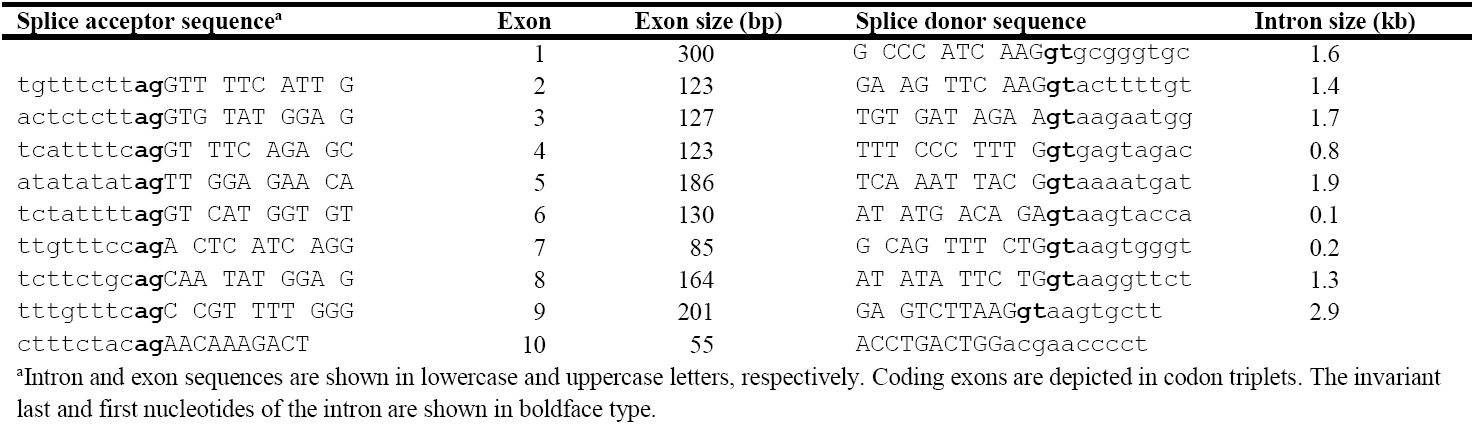
Exon-intron boundaries of mouse *Rbm45*

### Expression of human and mouse Rbm45 orthologues

Many RNA-binding proteins have been shown to be directly involved in tumorigenesis, including APOBEC-1 (Blanc *et al*., 2007), CUGBP2 (Mukhopadhyay *et al.*, 2003), Hel-N1, HuR, and musashi-1 (Sureban *et al.*, 2007, 2008). Moreover, CUGBP2, Hel-N1, HuR, and Musashi-1 are RDBPs similar in structure to Rbm45. Additionally, musashi-1 and Rbm45 are both expressed in differentiated neural cells and are involved in neural development (reviewed in Tamada *et al.*, 2002). Intriguingly, musashi-1 has been shown to be upregulated in colon cancer and knockdown of its expression inhibits tumorigenesis (Sureban *et al.*, 2008), properties expected of an oncogene (Hardin *et al*., 2012). Because of the similarities in both structure and expression of musashi-1 and Rbm45, coupled with the known property of cancer cells to have a dedifferentiated (embryonic-like) phenotype (Gilbert and Barresi, 2016), we hypothesized that expression of human *RBM45* would be dysregulated during colonic tumorigenesis. Consequently, qRT-PCR on seven paired samples of human normal and cancerous colon tissue was performed. Unlike musashi-1, no change in *RBM45* expression at the mRNA level was detected (*P* = 0.867, Fig. 3A); unfortunately, not enough tissue sample remained to determine protein levels. Had the resources been available, analyses of the expression of *RBM45* in developmental cancers, such as teratocarcinoma (tumor of stem cells; Gilbert and Barresi, 2016) and the two most common pediatric brain tumors, brain stem glioma and astrocytoma, two very aggressive cancers having a high rate of mortality and morbidity (Wrensch *et al*., 2002), may have illuminated a neural specific expression pathway with the potential to better understand how these tumors progress.

**Figure 3.**
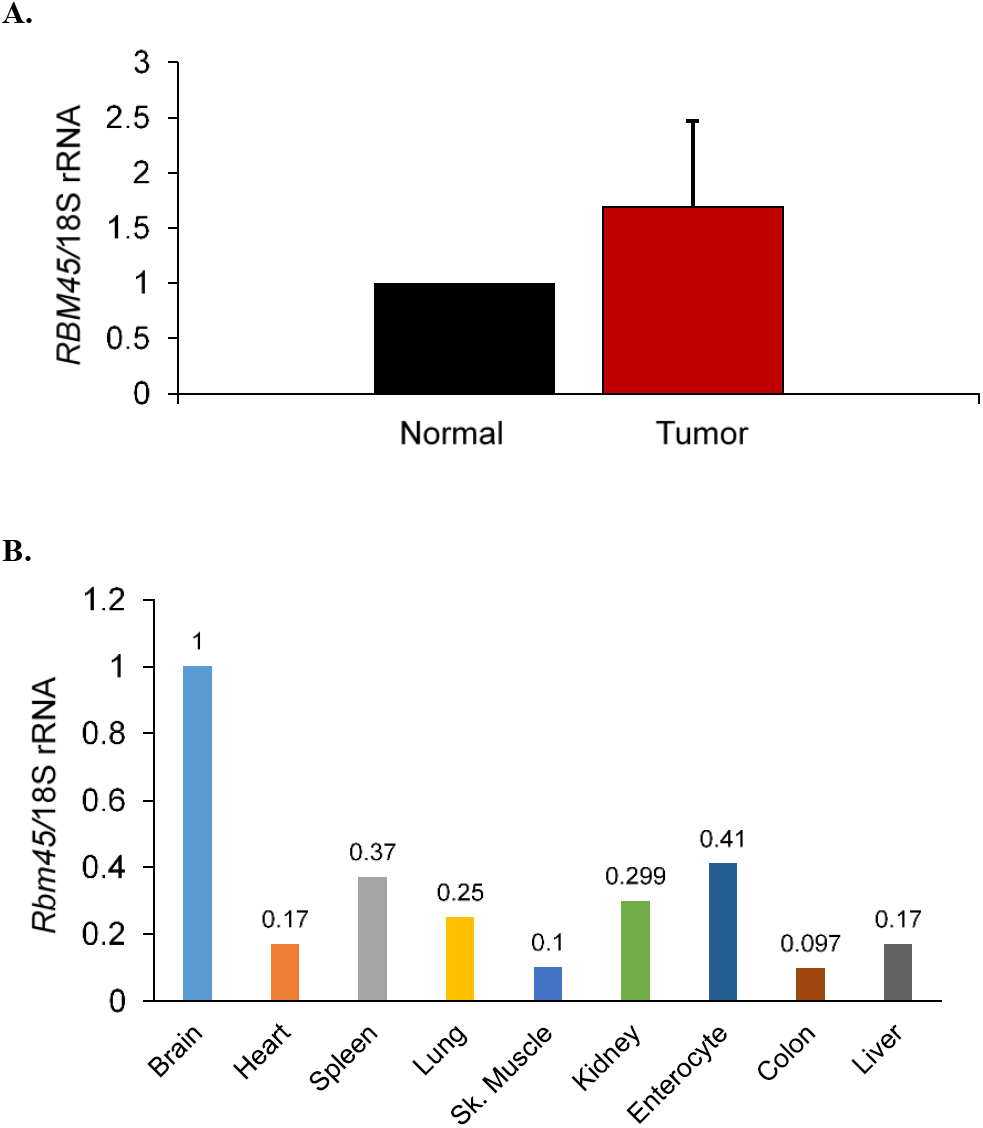
qRT-PCR of human *RBM45* / mouse *Rbm45* mRNA. (A) There is no significant difference in human *RBM45* expression in colonic tumors vs. matched adjacent normal tissue. The graph shows the ratio of human *RBM45* to the internal control, 18S rRNA, in human normal adjacent (Normal) and colorectal cancer (Tumor) tissue. *RBM45* / 18S rRNA levels in normal tissue were set to one. There is no significant difference in *RBM45* mRNA levels in tumor vs. normal tissue (*P* = 0.867, two-tailed *t* test; *n* = 7 matched normal and tumor samples). (B) Adult mouse brain tissue expresses the highest levels of *Rbm45* mRNA. Survey of adult mouse *Rbm45* mRNA tissue levels. *Rbm45* / 18S rRNA levels in brain were set to one. *Rbm*45 mRNA levels are reduced 10-fold in skeletal muscle (Sk. Muscle) and Colon and reduced 2.4-to 5.88-fold in Heart, Spleen, Lung, Kidney, Enterocyte (epithelial cells of the small intestine), and Liver. The 2 ^-ΔΔCt^ values are labelled above each tissue.

Concomitant with analysis of the tumor samples, we also examined the expression of mouse *Rbm45* in adult tissue including: brain, heart, spleen, lung, skeletal muscle, kidney, small intestinal enterocytes, colon, and liver. Our data (Fig. 3B) demonstrate that expression was highest in the brain with skeletal muscle and colon having 10-fold lower expression compared to brain; heart, spleen, lung, kidney, small intestinal enterocytes, and liver showed approximately three-to six-fold lower expression compared to brain. These data confirm and extend the work of Endo’s lab-group (2002) by directly quantifying steady-state tissue levels of mouse *Rbm45* mRNA.

### Recombinant expression and purification of human and mouse Rbm45

To begin the process of identifying potential RNA-binding partners, we cloned the human *RBM45* and mouse *Rbm45* cDNAs, respectively, to create Intein fusion proteins. The Intein moiety can be cleaved from the fusion protein during on-column purification to release the Rbm45 protein. Both human RBM45 and mouse Rbm45 were cloned so that the Intein protein is fused to the C-terminus.

Human RBM45 was efficiently expressed as a RBM45/C-terminal Intein fusion protein. Analysis of expressed RBM45-Intein protein by SDS-PAGE revealed a molecular mass of ∽80 kDa (53 kDa from RBM45 + 28 kDa from Intein), as expected. The recombinant protein was purified over a chitin column, subjected to on-column cleavage, and the eluent analyzed by SDS-PAGE revealing an ∽53 kDa protein at ∽95% purity (data not shown). The molecular mass of human RBM45 confirms that of Tamada and colleagues (2002). Having the ability to express large amounts of human RBM45 will allow us to identify putative RNA-(GACGAC motif; Ray *et al.*, 2013) and protein-binding partners in mouse brain extracts.

Expression of recombinant mouse Rbm45/C-terminal Intein fusion protein was poor. Expression optimization was carried out at 15°C, 25°C, or 30°C using a concentration gradient of 0.3, 0.4, or 0.5 mM of the inducer, IPTG. Only cells induced with 0.5 mM IPTG and incubated at 15°C produced detectable protein by SDS-PAGE analysis (data not shown). The fusion protein had the expected molecular mass of ∽80 kDa; however, expression was estimated to be 100-fold less than the control recombinant Intein-fusion protein, maltose binding protein. Furthermore, analysis of insoluble proteins in the pellet revealed no detectable Rbm45-Intein protein. Although expression of recombinant Rbm45 was low, we did not detect negative effects on bacterial replication in the presence of induced recombinant protein. Taken together, these data (not shown) indicate that mouse Rbm45 is not efficiently expressed from this vector. It is possible that mouse Rbm45 with Intein fused to the amino-terminus (i.e. N-terminus) could be expressed robustly in this system, but it is unclear why the highly related mouse and human Rbm45 proteins were not expressed similarly.

### Molecular Phylogenetics of Rbm45 Orthologues

In order to investigate the homology of Rbm45 across the tree of life, we searched the NCBI database for known and predicted orthologues. Our most recent search of the NCBI database (10/18/2017) for predicted Rbm45 orthologues revealed 214 gene loci. Similarly, a search of HomoloGene (https://www.ncbi.nlm.nih.gov/homologene/?term=RBM45; https://www.ncbi.nlm.nih.gov/gene/?Term=ortholog_gene_129831[group]), on the same date, showed orthologues across all metazoan taxa: animals that develop from only a blastula (one germ layer, no gastrulation), phylum Porifera (sponges); diploblasts (two germ layers), phylum Cnidaria (hydra); lophotrochozoan protostomia, phyla Mollusca (cephalopods: e.g. octopus), and Annelida (segmented worms); ecdysozoan protostomia, phylum Arthropoda (insects); and the deuterostomia phyla Echinodermata (sea urchin) and Chordata (Subphylum vertebrata: e.g. humans) (Hickman *et al*., 2011; see Brusca [2016] for an alternative view of the phylogeny). Our analysis of Rbm45 orthologues was confined to ten vertebrate species within classes Actinopterygii (ray-finned fishes), Amphibia, Reptilia (Aves), and Mammalia, including the important molecular genetic and developmental model systems *Danio rerio* (zebrafish), *Xenopus laevis* (African clawed frog), *Gallus gallus* (chicken), and *Mus musculus* (mouse). Unrooted phylogram analysis using Rbm45 cDNAs (Fig. 4A) demonstrate, as expected, the great apes (family Hominidae; human, chimpanzee, and orangutan) showing the closest evolutionary relatedness, with placentals as a monophyletic group, then a marsupial (opossum), followed by a reptile (chicken), amphibian (African clawed frog), and a fish (zebrafish). Pairwise alignment of Rbm45 nt sequences from human vs. chimpanzee and human vs. mouse revealed 97.8% and 90% identity, respectively (data not shown). Furthermore, multiple sequence alignment of Rbm45 orthologues from 10 different species within classes Mammalia, Reptilia (Aves), Amphibia, and Actinopterygii demonstrated 42.1% identity at the nt level (Fig. 4B, Supplementary Fig. 1), exceeding the threshold of identity due to weak selective pressure on degenerate codons (Koonin and Galperin, 2003).

**Figure 4.**
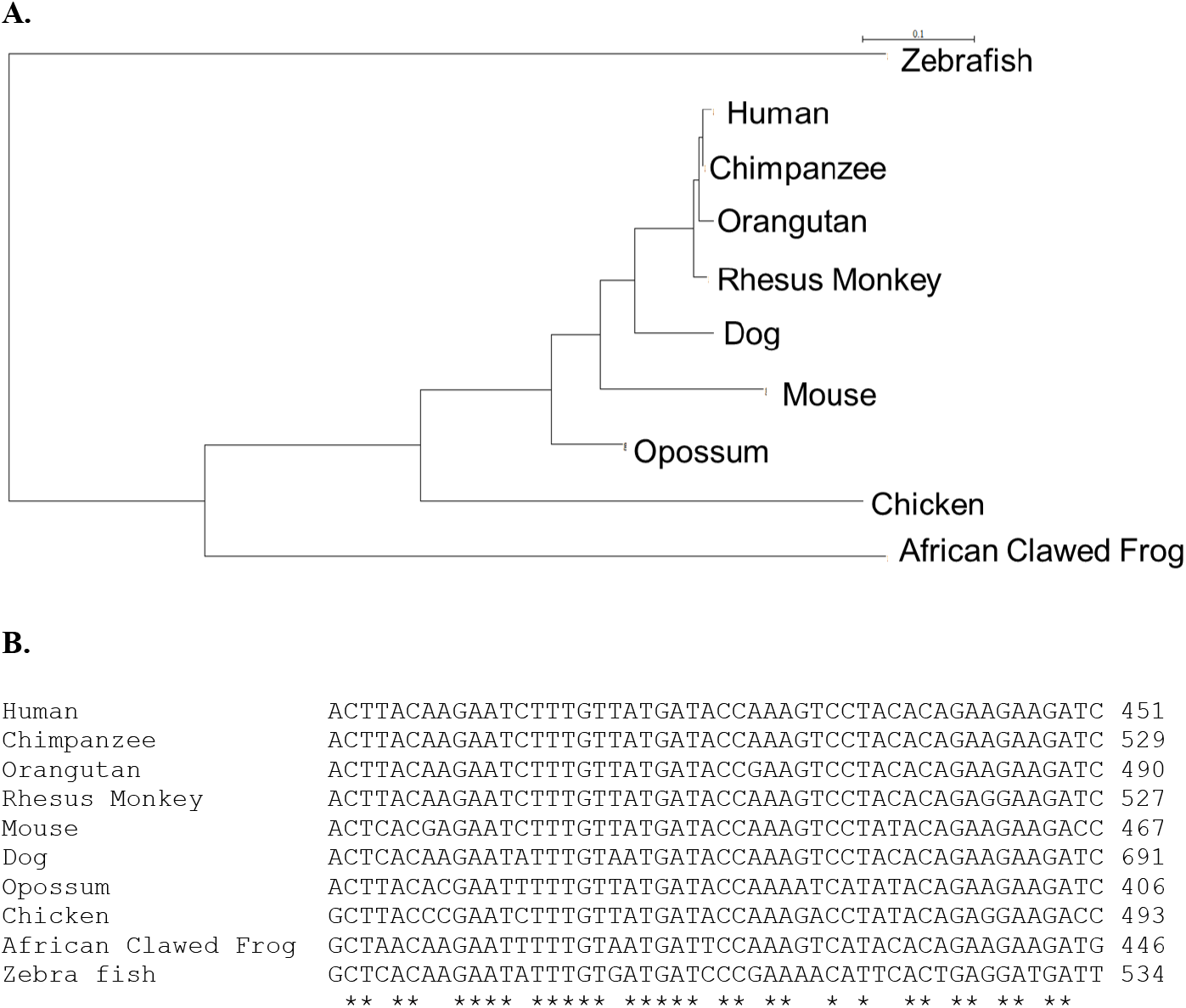
Molecular phylogenetics of Rbm45 cDNA sequences. Rbm45 nt sequence is highly conserved in vertebrates. (A) Phylogram created with the PhyML program using Rbm45 cDNA sequences from ten vertebrate species. The bar represents 0.1 nt substitutions per site. (B) Representative region of ClustalW multiple alignment of Rbm45 cDNA from ten vertebrate species. Asterisks (*) indicated identity. Species are listed with placentals first, then a marsupial, then egg-laying animals.

A similar examination using amino acid sequences from the same Rbm45 orthologues (Fig. 5) exhibit an almost identical phylogram (Fig. 5A) to that produced using nucleotide sequences (Fig. 4A), with the only difference being that mouse is not grouped with the other placentals (paraphyletic). However, other phylograms generated (not shown), illustrated the “correct” taxonomical relationships; we chose to use an unbiased methodology (see Experimental Procedures). Multiple sequence alignment of Rbm45 protein from 10 vertebrates demonstrates 65% conservation at the protein level (Fig. 5B, Supplementary Fig. 2). Since mouse is an important genetic model for understanding gene function in mammals, pairwise alignment of amino acid sequences from mouse and human was carried out revealing 93.5% conservation. In like manner, rat is often used as a physiological model for human metabolic pathways; pairwise alignment of mouse and rat Rbm45 showed 100% identity at the amino acid level (data not shown). Additionally, Tamada and coworkers (2002) reported 91.6% identity between rat and human Rbm45 protein. Unsurprisingly, orangutan, chimpanzee, and human, sharing a common ancestor (Darwin, 1874; Finstermeier *et al.*, 2013), form the least inclusive monophyletic grouping in the phylogram produced using Rbm45 amino acid sequence (Fig. 5A), with chimpanzee and human (subfamily Homininae, tribe Hominini [Gray, 1825; Hall and Hallgrimsson, 2014]) being the least inclusive monophyletic grouping in the phylogram generated using Rbm45 nt sequence (Fig. 4A).

**Figure 5.**
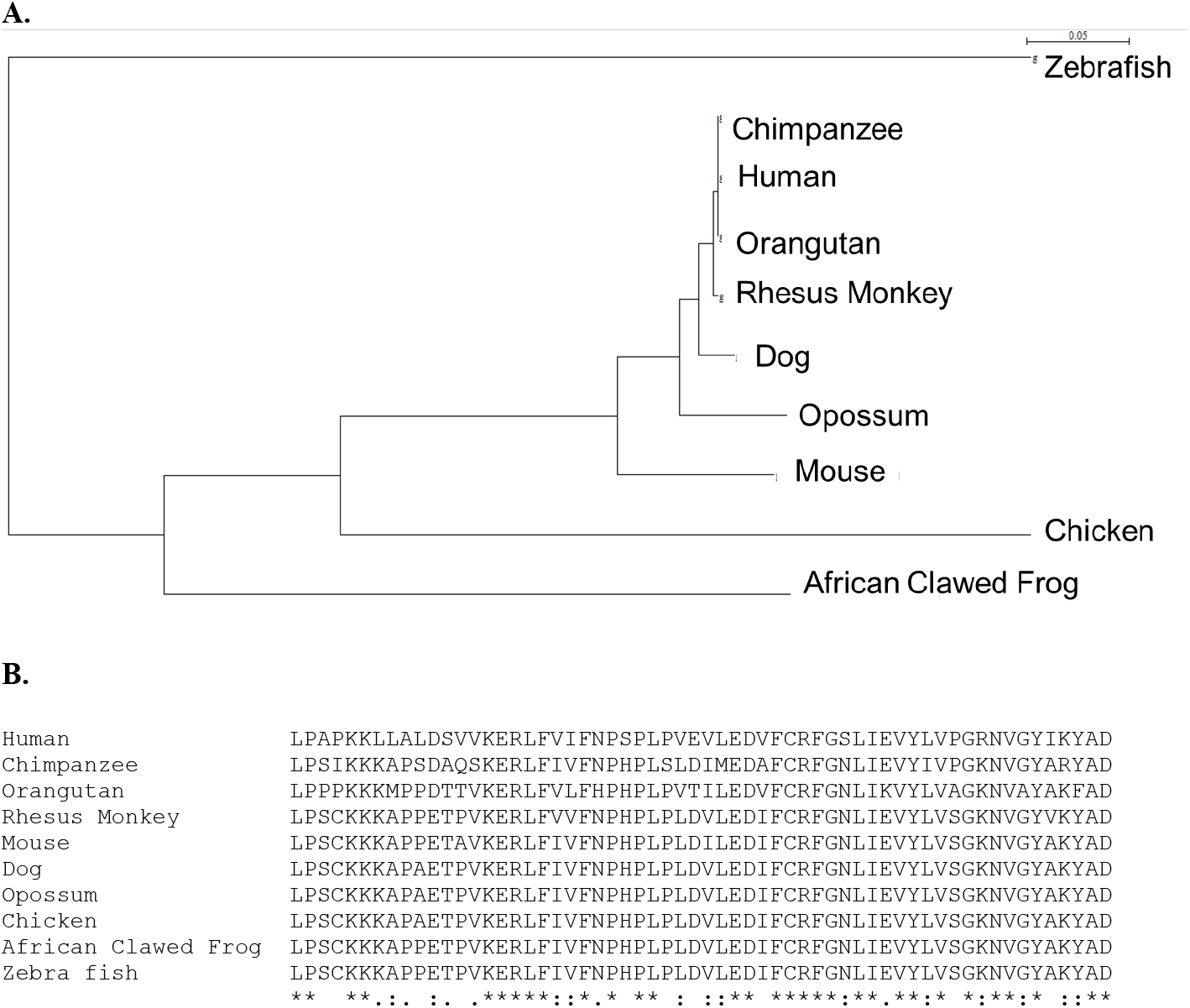
Molecular phylogenetics of Rbm45 protein sequences. Rbm45 AA sequence is highly conserved in vertebrates. (A) Phylogram created with the PhyML program using Rbm45 protein sequences from ten vertebrate species. The bar represents 0.05 AA substitutions per site. (B) Representative region of ClustalOmega multiple alignment of Rbm45 protein from ten vertebrate species. Asterisks (*) indicate identity, colons (:) indicate conserved AA change, and a single dot (.) indicates identity or conserved AA change in nine of the ten sequences. Species are listed with placentals first, then a marsupial, then egg-laying animals.

The phylograms in Figures 4A and 5A closely mimic the current understanding of tetrapod evolution (Amemiya *et al.*, 2013), supporting the molecular taxonomic relationship we observed. Considering our molecular phylogeny data and that sponges don’t have a nervous system, but retain a neural toolkit of genes (Leys, 2015), including an Rbm45 orthologue, it is tempting to speculate that these data suggest Rbm45 exhibits deep homology (Zimmer and Emlen, 2016), a fundamental, evolutionarily-conserved, function in (neuronal) development of metazoans.

## Acknowledgments

We thank Dr. Nicholas O. Davidson, Washington University School of Medicine, for tissue samples, reagents, hardware, and lab space for the qRT-PCR analysis on paired normal and cancerous human colon and mouse organ samples, Dr. Amy Greene for helpful comments on the manuscript, and Julie K. Henderson for editorial assistance. JOH thanks Dr. Shrikant Anant for recommending, in 2002, the newly discovered Rbm45 / Drbp1gene as a platform for undergraduate student research. This work was carried out from 2002 to 2017 and supported by a Ruth L. Kirschstein National Research Service Award T32 DK-07130 (2002-2003; JOH), Hope Scholars’ Grant from Tabor College (2008; JOH), science department funds from Trinity International University (2005-2006; JOH), Tabor College (2006-2008; ABLF, KMF, and JOH), Judson University (2012-2017; LEP, AV, KJL, AGE, DWH, and JOH), and a one-semester sabbatical leave from Judson University (2017; JOH).

